# Pathogen-inducible expression of autoactive NLRs confers multi-pathogen resistance in tomato

**DOI:** 10.1101/2024.08.30.610566

**Authors:** Wei Wei, Naio Koehler, Brandon Vega, Doogie Kim, Ashley Bendl, Buiry Min, China Lunde, Myeong-Je Cho, Ksenia Krasileva

## Abstract

Many disease resistance genes utilized in crop breeding encode nucleotide-binding leucine-rich repeat receptors (NLRs), which are frequently race-specific and fail to provide durable protection in the field. Consequently, engineering broad-spectrum resistance has become a primary goal. While autoactive NLRs can provide broad-spectrum immunity by bypassing specific pathogen recognition, they often impose severe fitness costs. Here, we hypothesized that precise transcriptional control could mitigate these penalties and developed a synthetic strategy coupling pathogen-inducible (PI) promoters with autoactive NLRs. Through a transcriptomic meta-analysis, we identified and validated native tomato PI promoters that respond to diverse pathogens while remaining insensitive to abiotic stress. We established a tunable system by pairing these promoters varying in basal and inducible expression with NLRs exhibiting different degrees of autoactivity. Systematic characterization revealed that balancing these components is critical to prevent leaky immune activation in the absence of pathogens. Transgenic tomatoes expressing a weakly autoactive NLR under a PI phenylalanine ammonia-lyase promoter exhibited enhanced resistance to two taxonomically distinct pathogens without growth impairment. Disruption of the PI sequence largely abolished this resistance, confirming the necessity of pathogen-mediated transcriptional induction. Finally, we added an additional layer of tunability by engineering the copy number of inducible elements and the core promoter to fine-tune promoter strength while preserving inducibility. Together, our results showcase a versatile framework for engineering multi-pathogen resistance and provide new insights into using transcriptional control in overcoming the fitness costs associated with autoactive NLRs.

## Introduction

Plant diseases contribute to significant crop yield losses worldwide, and disease resistance is the second most sought-after breeding characteristic after yield (Savary et al. 2019; Ristaino et al. 2021). Developing crop varieties with genetic resistance has been one of the most widely used strategies for controlling plant diseases (Chitwood-Brown et al. 2021; Liu et al. 2021; Lin et al. 2022; Khan et al. 2023), and reducing pesticide use. Many deployed resistance (R) genes to control biotrophic or hemi-biotrophic pathogens encode nucleotide-binding leucine-rich repeat (NLR) immune receptors, which recognize pathogen effectors and activate defense responses (Tamborski and Krasileva 2020; DeYoung and Innes 2006). However, NLR-mediated resistance is often limited in both spectrum and durability. Individual NLRs are typically activated by a single cognate effector present in only a subset of isolates from a given pathogen species, restricting their effectiveness against genetically diverse field pathogen populations (Van der Biezen and Jones 1998; Jones and Dangl 2006; Dodds and Rathjen 2010; Akamatsu et al. 2013). Moreover, because pathogens evolve rapidly, mutations or loss of the corresponding effector gene can enable evasion of NLR surveillance, leading to resistance breakdown (Stergiopoulos and de Wit 2009; Ortiz et al. 2022; Akamatsu et al. 2013; J. Chen et al. 2017). Thus, genetic resistance that offers broad-spectrum and durable protection is highly valuable and has been a long-standing goal for crop improvement.

Strategies to broaden the spectrum of resistance have been explored across diverse crop pathosystems, including stacking multiple resistance genes (Luo et al. 2021; Zhu et al. 2012), disabling host susceptibility genes (Fukuoka et al. 2009; Thomazella et al. 2021), and engineering NLR or pattern recognition receptor proteins (Kourelis et al. 2023; Yang et al. 2025; Seong et al. 2025). Constitutively active (autoactive or autoimmune) resistance genes represent an alternative approach that bypasses the requirement for specific pathogen recognition and trigger defense responses independently of effector perception, thereby conferring broad-spectrum resistance (Xiao et al. 2001; Clough et al. 2000; Li et al. 2001; Gao et al. 2011). Such autoactive R genes include truncated or mutated NLRs such as *RPW8*, *snc1* and the *RPM1^D505V^* allele (Xiao et al. 2001; Li et al. 2001; Gao et al. 2011), and non-NLR resistance genes such as wheat *Lr34*, which encodes an ABC transporter gene and Arabidopsis *dnd1* mutant which encodes a cyclic nucleotide-gated ion channel (Risk et al. 2013; Clough et al. 2000). However, constitutive activation of resistance is often accompanied by fitness costs, including dwarfism, reduced reproduction, and spontaneous lesions/necrosis (Clough et al. 2000; Xu et al. 2017; Boni et al. 2018). This growth-defense trade-off poses a major obstacle to the practical deployment of autoactive R genes in crop improvement.

Several studies have suggested that regulating the expression of autoactive R genes in a pathogen-inducible (PI) manner can substantially reduce fitness costs while maintaining resistance efficacy (Hutin et al. 2016; Xu et al. 2017; Gallas et al. 2024; Zeng et al. 2015; Boni et al. 2018). For example, expression of *Lr34* under the barley PI promoter *Hv-Ger4c* conferred robust resistance to powdery mildew and leaf rust without significant growth penalties (Boni et al. 2018). Several studies used the nature of pathogen transcription activator-like (TAL) effectors to induce host gene expression to regulate autoactive R genes (Hutin et al. 2016; Gallas et al. 2024; Zeng et al. 2015). Although effective, this strategy is limited to pathogens that deliver TAL effectors. A similar concept guides use of PI promoters to regulate a pathogen elicitor or an avirulence gene that activates an endogenous, non-autoactive NLR gene (Keller et al. 1999; Stuiver and Custers 2001; Honee et al. 1998; Kauder et al. 2025). Nevertheless, successful deployment of PI promoters to control autoactive R genes for broad-spectrum resistance still remains limited.

The success of this strategy relies heavily on the tight control of autoactive R gene expression. Such control requires not only strong induction during pathogen attack but also minimal expression under steady-state conditions, during other stresses, and across different tissues and developmental stages (Honee et al. 1998; Kauder et al. 2025). However, identifying promoters that respond exclusively to a single type of signal is challenging, as many genes, including defense-related genes, display pleiotropic expression patterns (Hou et al. 2022; Hendelman et al. 2021; Huang et al. 2010). For example, *pathogenesis-related Protein-1* (*PR-1*) genes have been used as a sensitive marker for defense responses, but they also largely respond to heat stress (Zaynab et al. 2021; Zribi et al. 2023). Controlled expression is further complicated by increasing evidence that many defense-associated genes are spatially regulated (Emonet et al. 2021; Munch et al. 2018; Beck et al. 2014). For example, the flagellin receptor gene *FLS2* is not only bacteria-inducible but also exhibits a specific spatial expression pattern in roots, with substantially higher expression in the endodermis than in meristematic cells (Beck et al. 2014). Despite the challenge of identifying pathogen-specific inducible promoters, the rapid expansion of RNA sequencing datasets over the past two decades provides a rich resource of gene expression patterns that could facilitate the discovery of novel pathogen-specific promoters (Tan et al. 2015; Knief 2014; Naidoo et al. 2018; Wei et al. 2022).

In this study, we developed a synthetic strategy using pathogen-inducible promoters to drive the expression of autoactive NLRs, conferring broad-spectrum resistance in tomato with minimal fitness costs. By leveraging extensive transcriptomic datasets, we identified candidate promoters that are responsive to diverse pathogens but remain insensitive to abiotic stress. Systematic evaluation revealed that precisely pairing promoter strength with NLR autoactivity is critical for preventing leaky defense activation. Our transgenic lines exhibited enhanced resistance to multiple pathogens without displaying growth defects under greenhouse conditions. Together, these results establish a flexible framework for harnessing transcriptional control to overcome the traditional trade-off between immunity and fitness.

## Results

### Selection of pathogen-inducible (PI) promoters by mining biotic and abiotic stress transcriptomic datasets

To achieve transcriptional control of autoactive NLRs that confers broad-spectrum resistance while minimizing fitness costs, it is necessary to identify PI promoters that respond broadly to pathogen infection but remain specific to pathogen-derived stimuli. We analyzed transcriptomic datasets of tomato leaves treated with four foliar pathogens from distinct taxonomic groups across kingdoms: bacteria *Pseudomonas syringae* pv tomato DC3000 (Pst) at 9 hour post inoculation (hpi) <u>(Yang et al. 2015)</u>, bacteria *Xanthomonas gardneri* (Xg) at 6 hpi (Thomazella et al. 2021), bacteria *Xanthomonas perforans* (Xp) at 48 hpi (Shi and Panthee 2020) and oomycete *Phytophthora infestans* (Pin) at 12 hpi (GSE33177). These four pathogens commonly cause diseases that affect tomato production, leading to severe yield loss (Jones et al. 2014). We also examined candidate promoters for their response to abiotic stressors by investigating transcriptomic datasets from tomato plants subject to drought stress (GSE39894), salt stress (GSE43492 and GSE16401) and wounding (GSE14637). In total, 98 genes showed significant expression induction after exposure to at least one pathogen (log_2_ fold change > 2 and FDR-corrected p-value < 0.05) but not during any examined abiotic stress (log_2_ fold change <1) (Fig. 1a). Although pathogens were chosen from distinct taxonomic groups, 23 genes were commonly activated by two or more pathogens and used for further analyses and validation (Fig. 1a; Fig. S1a; Table S1).

**Figure 1.**
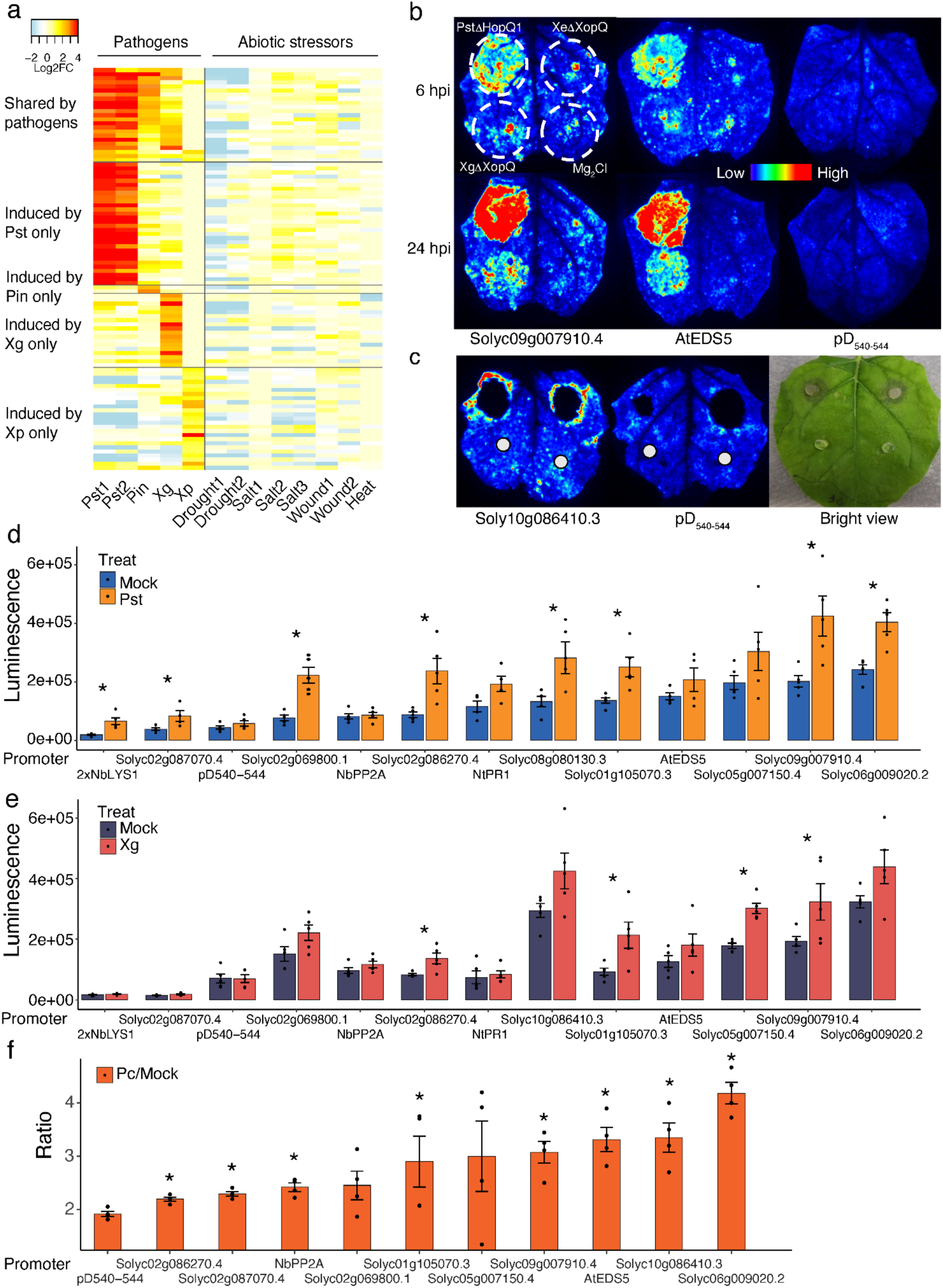
Selection and validation of PI promoters. **a,** The heatmap shows 98 genes that were significantly induced by at least one pathogen (log_2_ fold change > 2 and FDR-corrected p-value < 0.05) but not by abiotic stresses (log_2_ fold change <1). The color intensity represents the log_2_ fold change (log_2_FC) of the genes (row) under the treatments (column) in comparison with control treatments. The black horizontal lines in the heatmap separate different categories of genes. Pst, *Pseudomonas syringae* pv tomato DC3000; Pin, *Phytophthora infestans;* Xg, *Xanthomonas gardneri*; Xp, *Xanthomonas perforans*. **b,** Representative images of leaves expressing selected promoters, infiltrated with bacterial pathogens and the control. The whole leaf was first infiltrated with *Agrobacterium* carrying the promoter:luciferase constructs and two days later bacterial pathogens. The white dashed circles point to pathogen or MgCl_2_ infiltrated areas. Pst*ΔHopQ1*, Xg*ΔXopQ* and Xe*ΔXopQ* are the bacterial mutant strains with the corresponding effectors mutated to make virulent pathogens on *N. benthamiana* <u>(Schwartz et al. 2015; Schultink et al. 2017; Wei et al. 2007)</u>. Three promoters of genes known to respond to biotrophic pathogens or pathogen-associated molecular patterns (PAMPs) were included in the assay for comparison: *Arabidopsis* Enhanced Disease Susceptibility 5 (*AtEDS5*), *Nicotiana tabacum* Pathogenesis-Related 1 (*NtPR1*) and *Nicotiana benthamiana* lysozyme (*LYS1)* (Strompen, Grüner, and Pfitzner 1998; Nawrath et al. 2002; Garcia and Steinbrenner 2023). Promoters of two genes were used as negative controls: the *N. benthamiana* constitutive protein phosphatase 2A (*PP2A*) gene (Pombo et al. 2019) and the synthetic green-tissue specific gene promoter rice pD_540-544_ (Bai et al. 2020). **c,** Representative images of leaves expression selected promoters showed increased luciferase activities surrounding the necrotic lesion caused by *P. capsici*. The white circles indicate where the clean PDA plugs were put as mock inoculation. **d,e,** Quantification of luciferase activities driven by various PI promoters with and without PstΔHopQ1 (d), XgΔXopQ (e). Data are represented as mean ± standard error of 4-5 biological replicates. P-values were calculated by pairwise t-test and “*” indicates P<0.05. Multiple testing corrections using the Benjamini-Hochberg method. **e,** The ratio of luciferase activities of *P. capsici*-inoculated area by mock-inoculated area. The statistical comparisons were conducted between the negative control promoter pD_540-544_ and promoters of interest. P-values were calculated by pairwise t-test, *P<0.05.

To investigate which transcription factors (TFs) regulate these 23 common genes, we performed enrichment analysis of TF binding sequences in promoters and 5’UTRs (2.5kb upstream of the start codon). Twenty-two TFs were enriched and 17 of them belong to the MYB family, such as *SlMYB63* and *SlTHM27* that are known to be involved in defense response (Fig. S1b) (Fan et al. 2024; Pirona et al. 2023).

### Experimental validation of predicted PI promoters using reporter assays with *in vivo* pathogen challenge

An auto-luminescent luciferase-based reporter assay was used to validate the activity of predicted PI promoters via the *Agrobacterium*-mediated transient expression system on *Nicotiana benthamiana* (Garcia and Steinbrenner 2023). Fifteen out of the 23 promoters that were significantly induced by multiple pathogens were cloned successfully from tomato cultivar M82 (Table S1). After transient expression of these constructs in *N. benthamiana* leaves for 48 h, the infiltrated areas were challenged with the following pathogens to assess promoter responses to infection: Pst (*ΔhopQ1*), Xg (*ΔxopQ*), Xe (*ΔxopQ*), and *Phytophthora capsici* (Fig. 1b and 1c).

Eight promoters were significantly induced by Pst (*ΔhopQ1*) at 24 hpi, with luminescence differences between the pathogen- and mock-treated leaf areas ranging from 2.1- to 3.4-fold (Fig. 1d; Fig. S2). It is notable that these promoters exhibited different levels of basal expression without pathogen challenge, as seen in the mock-treated leaf areas (Fig. 1d). In addition, two positive control promoters, *AtEDS5* and *NtPR1*, which are known to be involved in basal defense (Strompen et al. 1998; Nawrath et al. 2002), showed some induction at 24 hpi but not high enough to reach statistical significance (Fig. 1d). Four out of 15 promoters displayed significantly induced responses to Xg (*ΔxopQ*) at 24 hpi, with fold differences from 1.6 to 2.3 (Fig. 1d; Fig. S2). Only one gene showed induction by Xe (*ΔxopQ*) (Fig. S2). After *P. capsici* inoculation, increased luciferase activity was observed at the leading edge of lesions at 40 hpi, where this hemi-biotrophic pathogen is posited to be in its biotrophic phase (Fig. 1c), and no luminescence was seen in the middle of lesions where cells were already dead. Because the negative control promoter *pD_540-544_* showed some background induction (Fig. 1c), promoter fold changes were compared to *pD_540-544_* to determine whether they were significantly induced by the pathogen. Eight promoters showed significant induction, with expression increases ranging from 2.2- to 4-fold after *P. capsici* infection (Fig. 1e).

Collectively, out of 15 promoters tested by luciferase assay, promoters of three genes were induced by all three pathogens: *Solyc02g086270.4* (a wall-associated kinase), *Solyc01g105070.3* (a peroxidase), and *Solyc09g007910.4* (a phenylalanine ammonia-lyase). Two additional promoters were induced by both Pst (*ΔhopQ1*) and *P. capsici*: *Solyc02g087070.4* (an alpha-dioxygenase) and *Solyc06g009020.2* (a glutathione S-transferase). These five promoters were selected for further analysis to regulate the autoactive NLRs.

### Deploying NLRs of different autoactivity levels with PI promoters of varying strength

As shown by the luciferase assays, some PI promoters exhibited high basal expression levels (Fig. 1c), which could lead to defense responses even in the absence of pathogens. To accommodate this variation in basal expression, we tested two NLRs with different levels of autoactivity: Sr33^E864V_V840M^ and Sr50, which originate from wheat and rye, respectively. In *N. benthamiana*, Sr33^E864V_V840M^, which carries two point mutations in Sr33, shows strong autoactivity while Sr50 exhibits inherently weak autoactivity (Tamborski et al. 2023). In principle, a strongly autoactive NLR should be driven by promoters with low basal expression, whereas a weakly autoactive NLR should be driven by promoters with higher basal expression (Fig. 2c). These pairings would likely minimize leaky activation of defense in the absence of pathogens while enabling robust defense upon induction.

**Figure 2.**
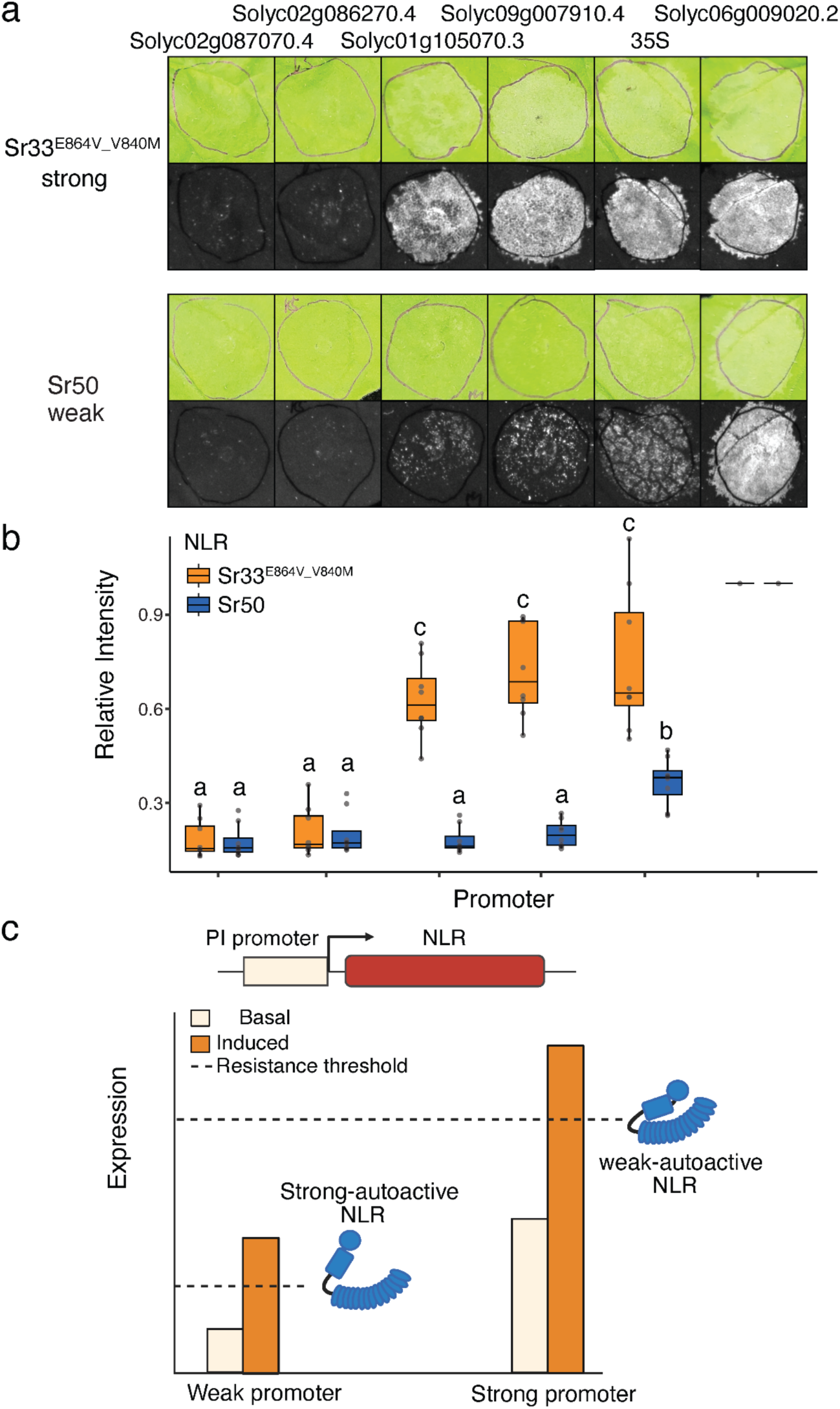
Deployment of NLRs with different autoactivities under the control of PI promoters. **a,** Two NLRs with different levels of autoactivity, driven by PI promoters with diverse basal expression, resulted in distinct levels of cell death without pathogen treatment. Constructs were transiently expressed in *N. benthamiana* leaves, and cell death was recorded two days after *Agrobacterium* infiltration. Images in the top row were taken under natural light. Images in the bottom row were acquired using excitation in the green visible spectrum with emission captured in the red visible spectrum, following an established method for visualizing leaf cell death <u>(Landeo Villanueva et al. 2021)</u>. **b,** Quantification of cell death based on emission light intensity. The promoter labels match those in the panel a. To account for variation in transient expression efficiency across leaves, light intensity was normalized to that driven by the *Solyc06g009020.2* promoter (the last column), a positive control which induced strong cell death under both NLRs. Statistical analysis was performed using two-way ANOVA with biological replicates (n=8), and different letters indicate significant differences after multiple testing corrections using the Benjamini-Hochberg method. **c,** Schematic illustrating the principles underlying the matching of PI promoters with NLRs of different autoactivities.

Combinations of the five selected promoters and two NLRs were transiently expressed in *N. benthamiana* without pathogen treatment. When Sr33^E864V_V840M^ was used, three promoters with higher basal expression, as well as the positive control promoter p35S, all caused widespread cell death, indicative of strong immune responses, whereas the two promoters with lower basal expression did not (Fig. 2a and 2b). In contrast, when Sr50 was used, two of the three strong promoters showed substantially less cell death, resulting in only sporadic lesions whose intensity were not significantly different from those using the two weak promoters with no visible cell death (Fig. 2a and 2b). The strongest promoter, *Solyc06g009020.2*, still induced widespread cell death but the positive control p35S resulted in moderate cell death, with an intensity lower than that driven by the strongest promoter, however higher than that driven by the other four promoters (Fig. 2a,b). Together, these results demonstrate that pairing PI promoters with different basal expression levels to NLRs with different autoactivities is an effective strategy to minimize leaky activation of defense in the absence of pathogens and to increase flexibility in PI promoter selection.

### A PI promoter combined with the weakly autoactive Sr50 confers enhanced resistance to multiple diseases in transgenic tomato without obvious fitness costs

Transgenic tomato plants expressing Sr50 with weak autoactivity regulated by the PI promoter *Solyc09g007910.4* were successfully generated and T_1_ plants with two copies of the transgene derived from three independent events were used for phenotyping. Tested lines showed significantly reduced bacterial growth after *Pst* at 4 days post inoculation (dpi), indicated by less bacterial fluorescence. They also showed significantly smaller lesion size after *P. capsici* inoculation at 4 dpi compared to inoculated wild-type tomato (Fig. 3a,b,c,d; Fig. S3), suggesting enhanced broad-spectrum resistance against taxonomically distinct pathogens. The acquired multi-pathogen resistance did not slow down vegetative growth or cause any obvious defects in reproduction in most transgenic events (Fig. 3e,f). In contrast, T_0_ plants expressing Sr50 regulated by the constitutive 35S promoter did not set fruit, and thus did not yield any T_1_ plants, suggesting that the use of PI transcriptional control may mitigate yield penalty.

**Figure 3.**
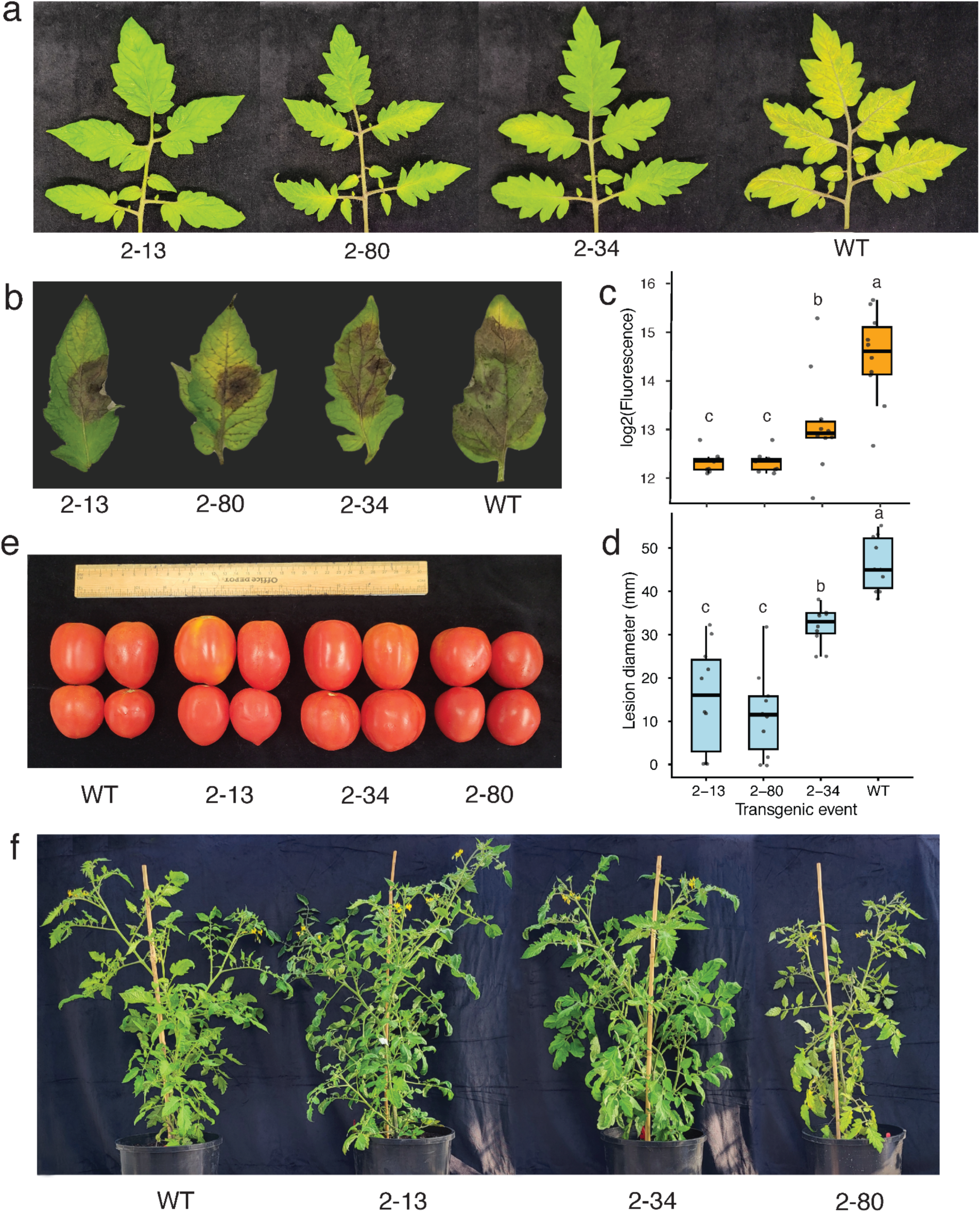
Transgenic tomatoes expressing Sr50 regulated by promoter *Solyc09g007910.4* led to multi-disease resistance with no obvious fitness cost. 2-13, 2-80 and 2-45, three independent transgenic events; WT, the wild-type, plants generated from the same batch of tissue culture but with no transgene insertion. **a,c,** The symptoms of transgenic tomato leaves after GFP-labeled *P. syringae* pv. tomato DC3000 inoculation. Symptoms and bacterial fluorescence were captured 4 dpi at OD_600_ 0.01. Statistical analysis was performed using non-parametric one-way ANOVA followed by pairwise Wilcoxon difference test, and different letters indicate significant differences (p<0.05) after multiple testing correction using the Benjamini-Hochberg method. Leaves (n = 10) were collected from two independent T_1_ plants, each carrying two transgene copies. **b,d** The symptoms of transgenic tomato leaves after *P. capsici* inoculation. Symptoms and lesion diameters were captured 4 dpi with a zoospore suspension density at 5 x 10^^3^/ml. Statistical analysis was described above. **e,** Representative fruits collected from transgenic tomatoes. **f,** Above-ground vegetative growth of transgenic tomato plants. Representative plants were photographed at 2.5 months of age.

### Truncation of the pathogen-inducible sequence largely reduces conferred resistance

To dissect promoter elements required for the resistance, we first identified the sequence responsible for pathogen inducibility in the promoter. The *proSolyc09g007910.4* was serially truncated from the 5’ end by ∼400bp increments and the truncated promoter variants were ligated to luciferase to evaluate expression (Fig. 4a). A significant induction by *PstΔHopQ1* disappeared when the sequence between - 1,408 bp and −1,027 bp was truncated, suggesting this region contains regulatory elements largely responsible for pathogen-inducibility (Fig. 4a). Transposase-accessible chromatin with sequencing (ATAC-Seq) data (Hendelman et al. 2021) showed that a high number of ATAC-seq reads mapped between −1,408 bp and −1,027 bp, supporting that this chromatin region is accessible for gene expression regulation (Fig. 4a). The 1.8 kb promoter was scanned with NewPlace and a total of 471 putative TF binding sites were identified (Supplementary Table S2), including multiple W-box and RAV1 TF binding sites, which are known to be involved in disease resistance and stress responses (Chandan et al. 2023; Javed and Gao 2023). Notably, a cluster of five W-box motifs were located within a region of 150 bp from −1,343 bp to −1,192 bp (Fig. 4a).

**Figure 4.**
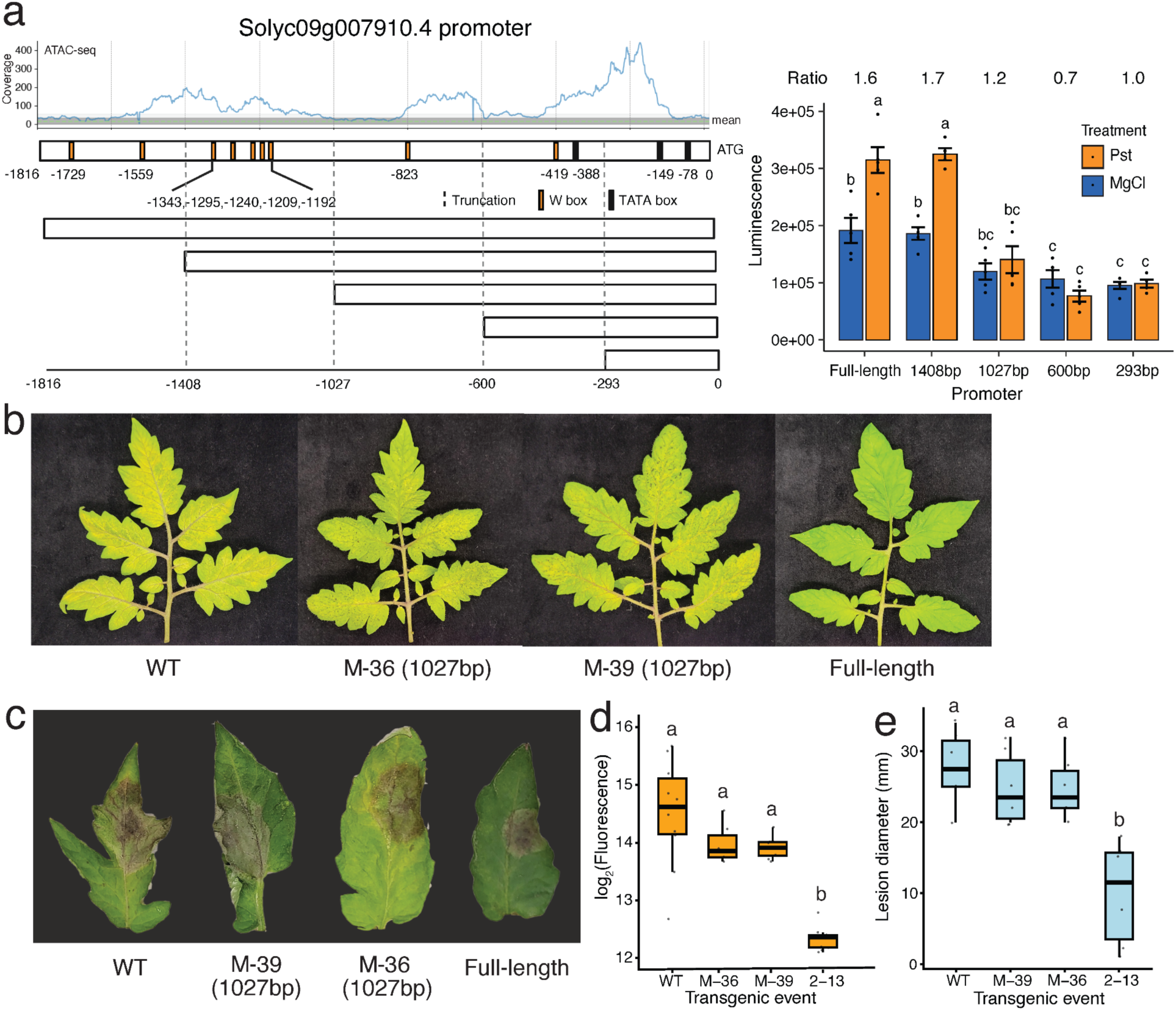
The mutant promoter with pathogen-inducible sequence removed largely reduces the resistance. **a,** Identification of pathogen-inducible sequences in promoter *Solyc09g007910.4* by 5’ end serial truncation. The top left shows the ATAC-seq read coverage plot of the promoter, with data from (Hendelman et al. 2021). The green dashed line indicates the genome-wide mean read coverage; the dark gray lines represent the mean ± 1 SD; and the light gray lines represent the mean ± 2 SD. The bottom left panel shows a schematic of the full-length promoter and truncation constructs. The bar plot on the right shows luciferase activity measured in *N. benthamiana* at 1 day post mock or *PstΔHopQ1* infiltration. Statistical significance was assessed using two-way ANOVA followed by pairwise comparisons (n = 4-5). Different letters indicate significant differences (p < 0.05) after Benjamini-Hochberg multiple testing correction. **b,d,** Disease symptoms of transgenic tomato leaves following inoculation with GFP-labeled *P. syringae* pv. tomato *DC3000*. M-36 and M-39 represent independent transgenic events carrying the promoter truncation variant; Full-length indicates transgenic tomato event 2-13 carrying the original promoter. See Fig. 3a for details of bacterial titer quantification and statistical analysis. **c,e,** Disease symptoms of transgenic tomato leaves following *P. capsici* inoculation. See Fig. 3b for details of pathogen quantification and statistical analysis.

Because pathogen inducibility was largely abolished in the −1027 bp to 0 bp promoter truncation variant (Fig. 4a), we generated transgenic tomatoes expressing Sr50 under the control of this variant promoter. These plants showed wild-type-like susceptibility to both *Pst* (Fig. 4b,e) and *P. capsici* (Fig. 4c,f), indicating that the PI sequence in the promoter is required for the enhanced resistance conferred by the full-length promoter.

### Another PI promoter combined with the strongly autoactive Sr33^E864V_V840M^ confers multi-disease resistance in leaf but impairs reproduction

We have also tested an additional PI promoter candidate in transgenic tomatoes. We generated transgenic T_0_ tomato plants expressing Sr33^E864V_V840M^ with strong autoactivity regulated by the PI promoter *Solyc02g086270.4*. T_0_ plants displayed vegetative growth comparable to that of their untransformed peer plants from the same tissue culture process (Fig. S4c), but none of them produced fruit (Fig. 4e). Upon closer observation, the stamens were enlarged and produced very little pollen, and the stigmas were thicker but shorter than those in wild-type flowers (Fig. 4e and Fig. S4a). *Solyc02g086270.4* is expressed approximately tenfold higher in flower tissues than in leaves (Fig. S4b), suggesting that higher expression in flower tissues may cause leaky immune responses in the development of reproductive organs, leading to anatomical changes and consequently impaired reproduction.

Due to the reproductive defect, each T_0_ was clonally propagated, and then inoculated with *P. syringae* pv. tomato DC3000. They had significantly less bacterial growth than the wild-type at 4 dpi, with greater than tenfold differences in bacterial titer (Fig. 5a,b). They also exhibited significantly smaller lesion sizes after *P. capsici* inoculation than did the wild-type tomato (Fig. 5c,d), indicating Sr33^E864V_V840M^ regulated by pro*Solyc02g086270.4* could confer effective resistance against multiple pathogens in leaves.

**Figure 5.**
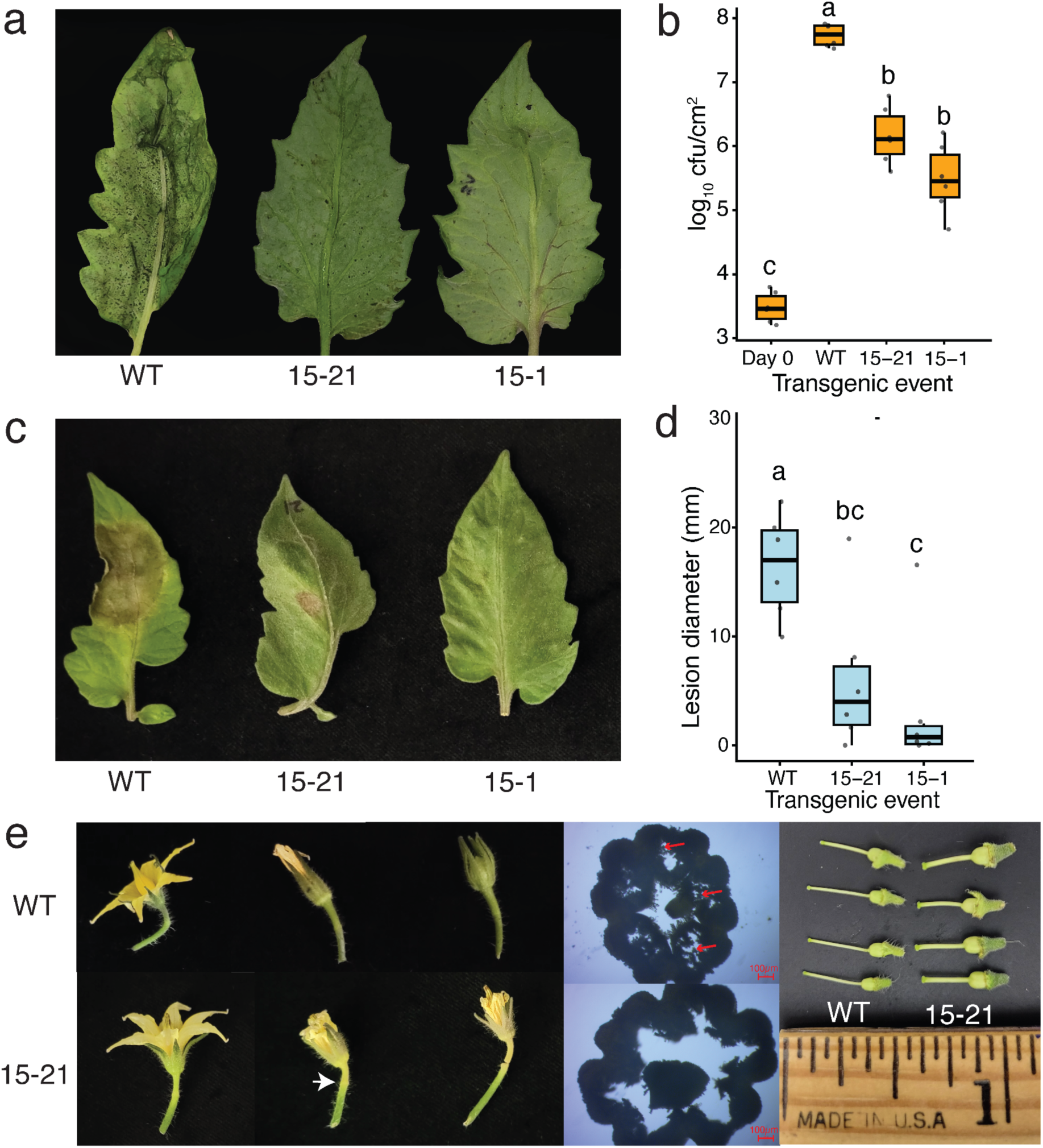
Transgenic tomatoes expressing Sr33^E864V_V840M^ regulated by promoter *Solyc02g086270.4* led to robust multi-disease resistance but impairs reproduction. 15-21, 15-1 and 15-3A, three independent transgenic events; WT, the wild-type, plants generated from the same batch of tissue culture but with no transgene insertion **a,b,** The symptoms of transgenic tomato leaves after *P. syringae* pv. tomato DC3000 inoculation. Symptoms and bacterial titer were captured four days post inoculation at OD_600_ 0.1. Statistical analysis was performed using non-parametric one-way ANOVA followed by pairwise Wilcoxon difference test, and different letters indicate significant differences (p<0.05) after multiple testing correction using the Benjamini-Hochberg method. Leaves (n = 6) were collected from two independent T_1_ transgenic plants generated by clonal propagation from T_0_ transgenics. **b,d,** The symptoms of transgenic tomato leaves after *P. capsici* inoculation. Symptoms and lesion diameters were captured three days post inoculation with a zoospore suspension density at 5 x 10^^3^/ml. Statistical analysis was described above. **c,** Comparison of flowers in T_0_ transgenic and wild-type plants. Flower-to-fruit set (left). White arrows indicate necrotic pedicels associated with flower abscission in all transgenic plants. Cross-sections of fused stamens showing pollen production (middle). Red arrows indicate pollen clusters in wild-type flowers, which are absent in transgenic flowers. Stigmas and ovaries are shown on the right.

**Figure 6.**
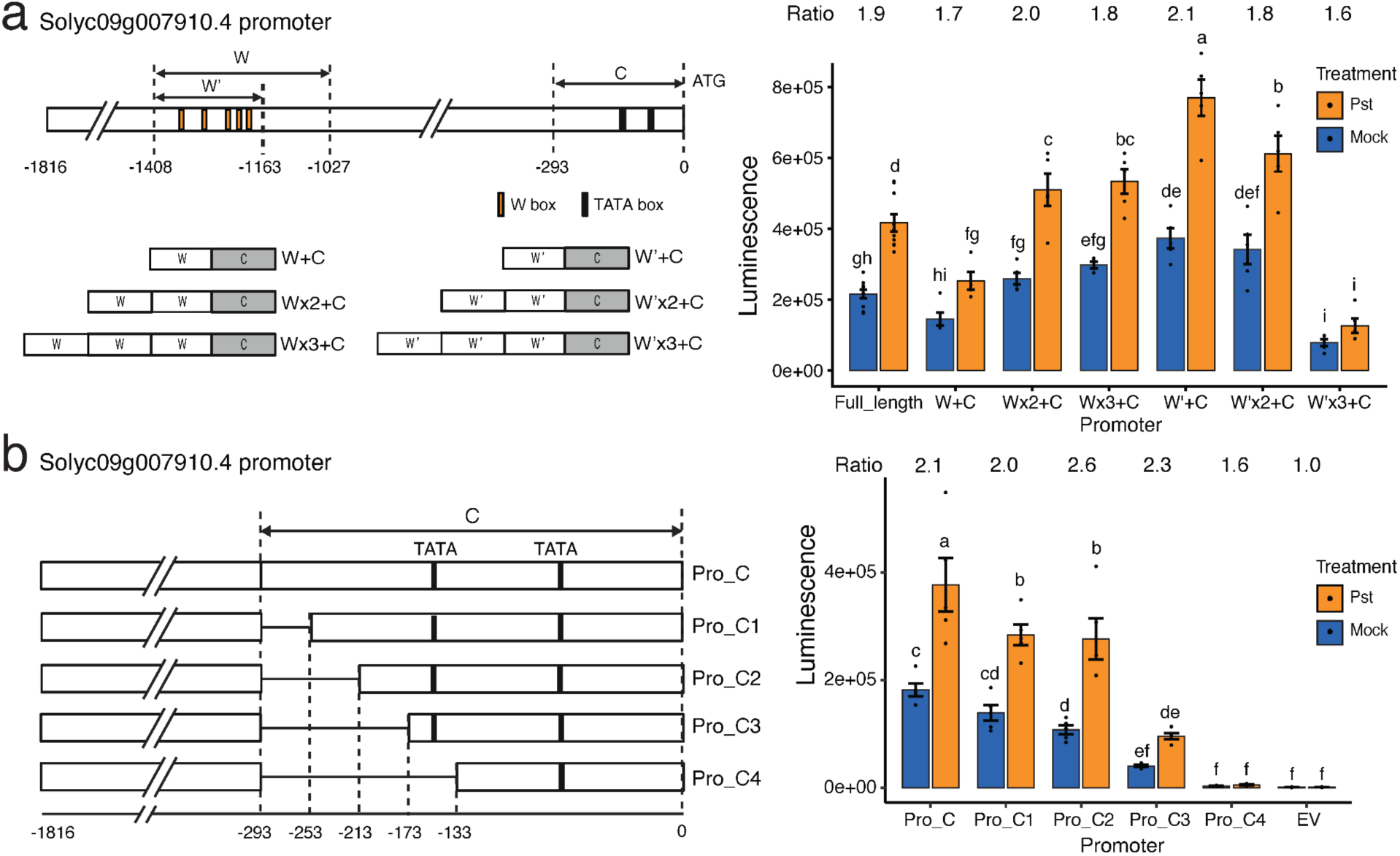
Promoter engineering to adjust basal expressions without affecting inducibility. **a,** Synthetic promoters with different repeat numbers of W sequence and W’ sequence. W, the ∼400bp pathogen-inducible sequences in pro*Solyc09g007910.4*. W’, the ∼250bp variant of the W sequence more specifically containing the five W-box motifs (see also Fig. 4a). Pst, *Pseudomonas syringae* pv tomato DC3000 *ΔhopQ1* mutant. **b,** Promoter with 5’ serial truncations of the core and proximal promoter. The luminescence was measured via a luciferase assay on *N. benthamiana* 1 day post PstΔHopQ1 and mock infiltration. Statistical analysis was performed using two-way ANOVA followed by pairwise difference test, with multiple testing p-value adjusted by the Benjamini-Hochberg method. Different letters indicate significant differences (p-value <0.05).

### Promoter engineering generated a gradient of expressions without affecting inducibility

Finally, we enhanced the tunability of our synthetic design through promoter engineering, focusing on whether basal expression levels could be modulated independently of pathogen inducibility. First, we generated recombined promoters for *proSolyc09g007910.4* by keeping only the PI sequence (W) from −1408 bp to −1027 bp and the non-PI region from −293 bp to 0 bp with TATA boxes, which we considered as the core and proximal promoter (C) (Fig. 4a; Fig. 5a). A variant of the W sequence (called W’) which is a shorter region (250 bp) that contains the five W-box motifs was examined in parallel (Fig. 4a; Fig. 5a). We tested whether adding different numbers of W or W’ affected expression. For W, one repeat yielded the lowest expression, and increasing the repeat number significantly increased both the basal and induced expression levels (Fig. 5a). The induction by *PstΔHopQ1* of different repeat numbers remained significant and were around 2-fold, which is similar to the full-length native promoter, suggesting that basal expression can be modulated without compromising inducibility. Interestingly, for W’, one repeat exhibited the highest basal and induced expression levels, and increasing the repeat number led to both lower basal and induced expression (Fig. 5a). With three repeats, the expression level was very low and the inducibility was nearly undetectable. This result suggests that the outcome of adjusting repeat numbers could be context- and sequence-dependent. We also applied the same engineering strategy to the promoter of *Solyc02g086270.4*. A region from - 500 bp to −400 bp was identified to be largely responsible for pathogen-inducibility, and adding three repeats increased both the basal and induced expression (Fig. S5a,b), further supporting the effectiveness of this strategy to fine-tune expression.

Second, we tested whether engineering the core and proximal promoter (C) could adjust the basal expression level. Serial 5’ truncations by 40 bp increments of the C in the full-length promoter resulted in gradual reductions in the basal expression levels, without affecting inducibility (Fig. 5b). The expression of Pro_C4, where the core promoter was truncated to 133 bp with the first TATA box removed, was reduced to the same level as the empty vector, indicating the essential role of primary transcription machinery binding elements in expression (Fig. 5b). This result supports that the strength of the core promoter could be rationally modulated. We also found the outcome of modulation was influenced by the sequence context. When linking core promoter variants with the 2x W sequence to make synthetic promoters, the gradual reduction in basal expression was not observed (Fig. S6). Overall these results showed that adjusting the repeat number of the PI sequence and core promoter can effectively modulate the basal expression without compromising the inducibility.

## Discussion

In this study, we present a framework for rational, pathogen-inducible (PI) promoter selection and synthetic biology design to confer multi-pathogen resistance in tomatoes. By mining transcriptomic data from tomatoes treated with various biotic and abiotic stresses, we selected PI promoters that were induced significantly by multiple pathogens but not responsive to investigate abiotic stress. Interestingly, many well-known defense-related genes, such as *PR1*, *EDS5* or the *WRKY70* (Han et al. 2023; Nawrath et al. 2002; Vidhyasekaran 2016), were not among our selected promoters due to their response to abiotic stressors. A comprehensive transcriptomic study showed that many genes rapidly upregulated by PAMPs (pathogen-associated molecular patterns) are also induced by abiotic stresses as part of a general stress response (Bjornson et al. 2021), likely reflecting extensive crosstalk among hormone signaling pathways (Robert-Seilaniantz et al. 2011; Shigenaga et al. 2017). In support of this, all thirteen PR1 genes in tomatoes were significantly upregulated by drought stress (Akbudak et al. 2020). Therefore, it is advisable to avoid using such genes for our purpose, especially with increasing extreme weather events through climate change, as deploying such genes responsive to abiotic stress might lead to fitness costs in regulating autoactive NLRs. Another interesting finding from our luciferase assay is that these conserved defense players showed less significant induction than our selected genes in susceptible host - virulent pathogen interaction. The expression of many known PI genes could be dampened by virulent pathogen effectors (Göhre et al. 2008; H. Chen et al. 2017). Overall, our result suggests that to identify PI promoters with desirable inducibility and specificity using a data-driven approach that incorporates susceptible hosts and different treatment conditions may be better than using empirical knowledge of conserved defense players.

We generated transgenic tomatoes that confer partial resistance to two pathogens from distinct taxonomic groups without displaying obvious growth defects, supporting the effectiveness of using PI promoters to regulate autoactive NLRs. Because native PI promoters differ in their basal expression level, an important insight from our study is that matching PI promoters with NLRs of appropriate autoactivity is critical for reducing leaky activation of defense in the absence of infection. The working construct in our transgenic tomatoes used the weakly autoactive NLR *Sr50*, highlighting the importance of tuning NLR sensitivity to balance resistance and fitness. Our findings align with previous studies showing that deploying a less sensitive resistance gene can mitigate fitness costs. For example, *Lr34*, used to confer both powdery mildew and rust resistance in barley, encodes a non-NLR protein that provides quantitative, partial resistance and reduces disease severity without fully stopping disease progression <u>(Risk et al. 2013; Boni et al. 2018)</u>. In another example, co-expression of the NLR gene *R3a* and its cognate effector *Avr3a* under the control of a PI promoter showed that using an *Avr3a* allele with reduced cell-death–inducing activity alleviated the growth impairment observed with other alleles (Kauder et al. 2025). In conclusion, these findings suggest that future efforts should combine refined promoter selection with protein engineering. Such a dual approach is now increasingly practical, leveraging recent breakthroughs in the NLR biology and molecular dynamics of NLR-AVR interactions <u>(Tamborski et al. 2023; Seong et al. 2025)</u>.

In our study, transgenic tomatoes expressing the strongly autoactive *Sr33^E864V_V840M^* under the PI promoter *Solyc02g086270.4* exhibited strong resistance, but the plants did not set fruit. This phenotype is most likely due to elevated promoter activity in reproductive tissues. Notably, although the strong autoactive construct caused reproductive defects, this feature could be advantageous in specific applications. For example, an embryo-lethal cassette may be useful in crop commercialization strategies, particularly in clonally propagated crops or vegetative crops where disease resistance and yield can be uncoupled from sexual reproduction. *Solyc02g086270.4* encodes a protein from the wall-associated kinase (WAK) family involved in perceiving damage-associated signals under pathogen attack. Expression of WAKs can be induced upon pathogen attack but was also detected in various organs and developmental stages, reflecting additional roles in plant development (Sun et al. 2020; Kurt et al. 2020). Like WAKs, many native PI genes harbor tissue-specific expression patterns (Zribi et al. 2023; Shi et al. 2024) and many others even exhibit spatial regulation at cell-type resolution (Beck et al. 2014; Nobori et al. 2025). Together, these observations suggest that achieving broad-spectrum resistance without fitness penalties may require an additional regulatory layer to refine spatial and developmental specificity. Incorporating synthetic regulatory circuits could allow immune activation to be confined to pathogen-threatened tissues while avoiding sensitive stages such as reproduction (Brophy et al. 2022; Khan et al. 2025).

Engineering PI promoters could represent an effective complementary strategy for fine tuning the developmental system to balance immunity and growth. In this study, we generated PI promoter variants displaying gradient basal expressions while maintaining pathogen inducibility. To modulate basal expression, we explored multiple engineering strategies, as it was shown that promoter activity could be affected by number, order, spacing and position of cis-elements and core promoters (Rushton et al. 2002; Kauder et al. 2025; Huang et al. 2025; Cai et al. 2020). However, generalizable engineering rules remain largely elusive, especially for inducible promoters that have both basal and inducible expression levels. Manipulating the copy number of inducible elements revealed context-dependent effects: increasing the motif number either elevated or reduced basal expression depending on whether the W or W′ element was used. Although both W and W′ contain W-box clusters, W′ is a truncated version of W, and their contrasting behaviors may reflect differences in spacing among W-boxes or between W-box clusters and the core promoter. Positional effects were also observed when modifying the core promoter. Core promoter variants displayed distinct basal expression levels in the native promoter context, yet these differences were attenuated when positioned adjacent to inducible elements in synthetic constructs. Similar positional effects of cis-regulatory elements have been reported in other systems, although the underlying mechanisms remain unclear <u>(Murphy et al. 2007; Jores et al. 2024; Voichek et al. 2024)</u>. The complexity of promoter architecture and the dynamic nature of the cis-regulatory grammar highlight the need for large-scale, systematic exploration to achieve predictable PI promoter engineering.

In conclusion, our study integrated transcriptome meta-analysis with synthetic biology to engineer multi-pathogen resistance without obvious growth penalties, thereby providing a valuable resource for crop improvement. Although not directly tested here, the synthetic resistance strategy developed may offer enhanced durability compared with traditional NLR-based resistance, as it bypasses specific NLR-effector recognition and thus may reduce the likelihood of pathogen evasion. We used tomato as a proof of concept due to its economic importance and amenability to transformation; however, the framework described here is generalizable. Many core signaling components of NLR-mediated immunity are conserved across plant species, and it is now easy to identify PI promoters with the abundant transcriptomic data (Maekawa et al. 2012; Dongus and Parker 2021; Ramírez-Zavaleta et al. 2022). This approach may be particularly valuable for diseases lacking effective genetic resistance resources, such as Huanglongbing in citrus (Alves et al. 2020), and other emerging pathogen threats.

## Materials and Methods

### Plant growth

*Nicotiana benthamiana* plants were grown in a growth chamber (Conviron) under 25 °C/23 °C, a 14 h/10 h day/night cycle and 80 µmol m^-2^ s^-1^. The seeds were sowed on the soil surface and covered with a plastic dome to increase humidity until germination. The seedlings were transferred to 3-inch pots 10 days post sowing. Tomatoes were grown in a greenhouse with the condition 28 °C/22 °C, 14 h/10 h day/night cycle. Fertilizers for tomato include Calcinit (YaraLiva) and 20-20-20 General Purpose (Peters Professional), supplemented with magnesium sulfate (Giles) and spraying Calcium chloride (1.6%) on to foliage and early developing fruits.

### PI promoter selection

PI promoters were selected using the GeneVestigator (Immunai) that enables cross-comparison between transcriptomic datasets generated with different platforms (RNA-seq or Microarray) and different versions of tomato reference genomes. The promoters of differentially expressed genes (DEGs) after pathogen infection (log_2_ fold change >2 and P value<0.05) were selected and filtered by no significant induction by abiotic stress treatment (log_2_FC<1).

### Plasmid construction

Constructs in this study were made through GoldenGate cloning or Gibson assembly cloning methods if relevant restriction enzyme sites were prevalent in the sequence (Engler and Marillonnet 2014; Gibson et al. 2009). DNA parts including 35S promoter, 6xHA tag and heat shock protein 18.2 (HSP) terminator were obtained from the Moclo Plant Toolkit (Engler et al 2014). PI promoters were cloned from the tomato genomic DNA using the Phusion DNA polymerase (New England Biolabs, USA) with addition of overhangs and the BsaI restriction enzyme site for downstream reactions. Primers were designed based on the *Solanum lycopersicum* ITAG4.0 genome targeting the 2kb∼3kb genomic regions upstream of the start codon of the gene, including promoter and 5’UTR (Supplementary Table S1). For luciferase assay, promoters were ligated via GoldenGate cloning into the FBP vector (kindly given by Dr. Steinbrenner’s lab) <u>(Garcia and Steinbrenner 2023)</u>. Synthetic promoters were made by cloning part sequences from native promoters and stitching them together using GoldenGate cloning. For stable transformation constructs, coding region of Sr33^E864V_V840M^ and genomic DNA of Sr50 containing 5’UTR, coding sequences, introns and 3’UTRs were ligated with PI promoters or synthetic promoters and cloned into pCAMBIA2300 vector with a kanamycin selection marker (Tamborski et al. 2023; Simon et al. 2021).

### *Agrobacterium*-mediated transient expression

*Agrobacterium tumefaciens* GV3101:pMP90 transformed with the corresponding construct was grown at 28 °C in Luria-Bertani (LB) medium containing with 50 mg ml^-1^ rifampicin, 25 mg ml^-1^ gentamycin and either 100 mg ml^-1^ carbenicillin or 50 mg m^-1^l kanamycin depending on the vector. The cultures were centrifuged at 4,500 g for 5 min and the pellet resuspended in infiltration medium (10 mM MES pH 5.6, 10 mM MgCl2 and 150 µM acetosyringone). Optical density was adjusted to a final optical density at 600 nm (OD_600_) 0.1 as this concentration of *Agrobacterium* was tested to achieve optimized expression efficiency and single-copy insertion frequency in transient assays (Carlson et al 2023). For co-infiltrations, the suspensions of individual constructs were mixed in a 1:1 ratio at a final density of 0.1 for each strain. The top two fully expanded leaves of 4-5 weeks old *N. benthamiana* plants were infiltrated using a blunt syringe. Plants were transferred to dim light (∼30 µmol m^-2^ s^-1^) after infiltration for better transient expression (Jutras et al 2021). Phenotypes were recorded 2-3 days after infiltration. For luciferase assays, we noticed that the luciferin travels outside the *Agrobacterium* infiltration area and causes cross-contamination so we only infiltrated two spots on each side of the main leaf vein.

### *N. benthamiana* promoter luciferase assay

Two days post *Agrobacterium* infiltration, leaf areas transiently expressing luciferase were inoculated with pathogens to test the inducibility. Since the wild-type tomato bacterial pathogens in the transcriptomic analyses are not compatible with *N. benthamiana*, we used the compatible mutants which have specific effector knocked-out. The following pathogen mutants were used: the *Pseudomonas syringae* pv tomato DC3000 *hopQ1* knockout mutant (Pst*ΔhopQ1*), the *Xanthomonas gardneri xopQ* knockout mutant (Xg*ΔxopQ*), and the *Xanthomonas euvesicatoria xopQ* knockout mutant (Xe*ΔxopQ*) were obtained from Dr. Brian Staskawicz’s lab (Schwartz et al. 2015; Schultink et al. 2017; Wei et al. 2007). Due to lack of a compatible *X. perforans* strain, the Xe*ΔxopQ* was used as a substitute as studies showed they are very closely related (Jibrin et al 2018). Bacterial strains were grown on a LB medium plate containing 100 mg ml^-1^ rifampicin at 28 °C for two days. On the night before inoculation, a pellet of bacteria was inoculated into liquid NYG (0.5% peptone, 0.3% yeast extract, 2% glycerol) with 100mg ml^-1^ rifampicin and cultured for 16 h at 28°C. On the next day, bacteria cells were harvested by centrifuging at 4,500 x g for 5 min and resuspended in 10 mM MgCl_2_ with OD_600_ adjusted to be 0.02 for Pst*ΔhopQ1* and 0.25 for Xg*ΔxopQ* and Xe*ΔxopQ* (Xu et al. 2017; Thomazella et al. 2021). Bacteria solution was infiltrated into the same infiltration spot as *Agrobacterium*, with a syringe to make a puncture first to enhance infiltration. The *Phytophthora capsici* strain MP24 was obtained from Dr. Matteo Garbelotto’s lab, due to the incompatibility of *N. benthamiana* and *P. infestans*. After growing on the potato dextrose agar (PDA) for 5-7 days, mycelial discs were punched using a 6 mm cork borer on the edge of the colony and put on the adaxial side of detached *N. benthamiana* leaves. Mock treatments used PDA punches without *P. capsici*. Detached *N. benthamiana* leaves were put in a humid incubation box covered with wet paper towels.

After 6 hpi and 24 hpi of bacteria infiltration and 40 hpi of *P. capsica* inoculation, luminescence was imaged with a ChemiDoc MP imaging system (Bio-Rad, USA) using the “Chemiluminescence” function with a 60s exposure time. For quantification, the luminescence was measured using an Infinite F Plex Plate Reader (Tecan, Switzerland). Three leaf discs were measured from the same infiltration area as technical replicates and five biological replicates were used. For *P. capsica* quantification, the mean color intensity of the peripheral area surrounding the necrotrophic lesion was measured using the software ImageJ <u>(Schneider et al. 2012)</u> and compared to background intensity.

### Tomato transformation and transgenic plants screening

Tomato transformation was conducted in the cultivar M82 using *Agrobacterium* strain AGL1 at the Plant Genomics and Transformation Facility at the Innovative Genomics Institute (IGI Berkeley). T_0_ plants derived from tissue culture were genotyped for presence of transgene with the Phire Plant Direct PCR Master Mix (Thermo Scientific). For transformants that can produce fruits, transgene copy number of T_1_ plants was determined by DNA extraction using the CTAB method (https://opsdiagnostics.com/notes/protocols/ctab_protocol_for_plants.html) and a combination of droplet digital PCR with the QX200™ ddPCR™ EvaGreen Supermix on a QX200 system (Głowacka et al. 2016) and quantitative PCR using PowerUp SYBR Green Master Mix (Applied Biosystems) on a QuantStudio™ 5 Real-Time PCR System.

For transformants that cannot produce fruits, tomato clonal propagation was performed. Young growing shoots with 2-3 expanded leaves were cut. The cutting interface was dipped with Rooting Powder (Bonide Bontone II, Amazon) and planted in soil with 1:2:2 Supersoil:perlite:vermiculite. Transplants were grown in a shaded area for three weeks until new roots grew substantially. The clonally propagated plants were genotyped to confirm the presence of transgene.

### Tomato disease phenotyping assay

The tomato inoculation with *P. syringae* pv tomato DC3000 were performed mostly following the protocol described previously (Thomazella et al. 2021). Briefly, bacteria were grown in NYG liquid media for 16 h at 28 °C. Cells were resuspended in 10 mM MgCl_2_ and Silwet L-77 at OD_600_ 0.1 or 0.01 (see figure legends). Detached plant leaflets were dipped into the cell suspension for ten times and were subsequently put in a humid incubation box covered with wet paper towels at 25 °C and 150 µmol m^-2^ s^-1^ until disease symptoms developed. For bacterial quantification, three 8 mm leaf disks from one leaf were punched using cork borers, ground in 300 µL 10 mM MgCl_2_ in a 2 ml Eppendorf tube with two 3.2 mm stainless beads using a Biospec mini-beadbeater (1500 rpm, vital distance 3.175 cm). For quantification of GFP-labeled *P. syringae* pv tomato DC3000, 50 µL of homogenized leaf extracts was put in a 96-well plate (Greiner, Black) and measured the GFP fluorescence using an Infinite F Plex Plate Reader (Tecan). Three technical replicates were measured from the leaf extract. If the resistance was too strong so that the fluorescence did not reach the detection threshold, a colony forming unit was counted. More specifically, 10x serial dilutions of the homogenate were plated onto LB agar supplemented with 100 μg/ml rifampicin and 50 μg/ml cycloheximide for selecting *P. syringae* pv tomato DC3000 and suppressing fungal growth. Colonies were observed and counted 72 h after plating. The method to induce *P. capsici* zoospore was previously described in <u>(Kaur et al. 2024)</u>. The isolates were grown on clarified V8 agar media (https://wiki.bugwood.org/Clarified_V8_agar). The *P. capsici* isolates MP24 was grown on a plate for 7 days at room temperature in the dark. Then, 5 cm plugs were excised from these Petri dishes using a cork borer. A total of 5-7 of these plugs were placed in deep petri dishes with 25 mL of sterile distilled water and placed in an incubator at 25 °C for 48 h with fluorescent lighting. Zoospore release was induced by transferring these dishes to a 4 °C cold chamber for 60 min and then placing them back in the incubator for 30 min. A cheesecloth was used to filter the contents of the deep petri dishes to remove agar plugs. The resulting solution was subjected to zoospore quantification using a hemocytometer (Hausser Scientific, PA, USA). The final concentration of zoospores was adjusted using sterile distilled water to 5 x 10^3^/ml. Leaves were spot inoculated by pipetting 10 µl droplets of the spore suspension on the adaxial side of each tomato leaflet.

## Acknowledgements

We thank Dr. Adam Steinbrenner (University of Washington) for providing the luciferase construct. We are grateful to Tina Popenuck and Dr. Matteo Garbelotto (UC Berkeley) for the *Phytophthora capsici* strain. We also thank Douglas Dahlbeck and Dr. Brian Staskawicz (UC Berkeley) for the *Pseudomonas syringae* pv. tomato DC3000 *HopQ1* knockout mutant, the *Xanthomonas gardneri XopQ* knockout mutant, and the *Xanthomonas euvesicatoria XopQ* knockout mutant. We appreciate Christina M. Wistrom (UC Berkeley) for greenhouse management and plant care. We thank Dr. Sheila McCormick for guidance on flower dissection and anatomical analysis and review of the manuscript. Finally, we acknowledge members of the Krasileva lab, Dr. Rakesh Kumar and Dr. Lorena Parra for helpful discussions, and Dr. Kyungyong Seong, Dr. Rakesh Kumar, and Chandler Sutherland for manuscript editing.

## Author contribution

W.W. and K.K. conceptualized this study and designed the experimental plan. W.W. performed the experiments, data analysis and wrote the manuscript. B.V., D.K., A.B and C.L. assisted in experiments including promoter cloning and engineering, promoter luciferase assay, planting, and transgenic genotyping and phenotyping. N.K. and B.M. conducted the tomato transformation supported by M.-J.C.

