## Supplementary_figures for "Pathogen-inducible expression of autoactive NLRs confers multi-pathogen resistance in tomato"

a

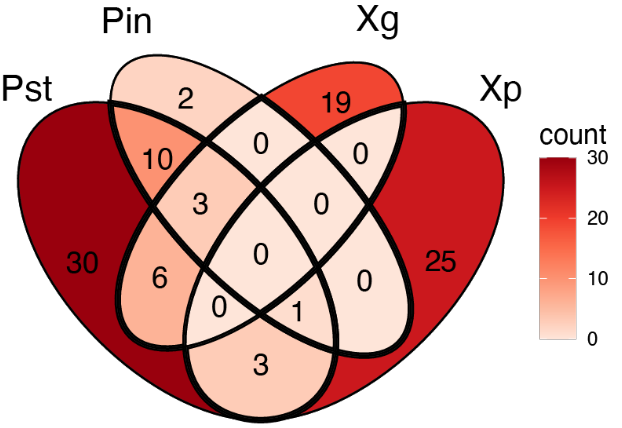


b

| TF | Gene ID (ITAGv2.4) | Motif logo | Fold enrichment | GO terms of ortholog in Arabidopsis |
|---|---|---|---|---|
| SlMYB17 | Solyc01g094360.2 | 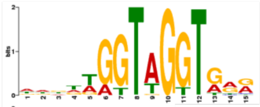 | 4.6 | GO:0009751<br>response to salicylic acid<br>GO:0009753<br>response to jasmonic acid |
| SlMYB93 | Solyc04g056310.2 | 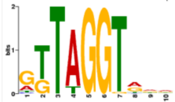 | 7.6 | GO:0009751<br>response to salicylic acid<br>GO:0009753<br>response to jasmonic acid |
| SlMYB63 | Solyc10g005550.1 | 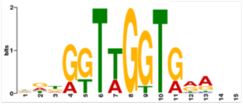 | 5.4 | GO:0009809<br>lignin biosynthetic process<br>GO:2000652<br>regulation of secondary cell wall biogenesis |
| SlTHM27 | Solyc10g055410.1 | 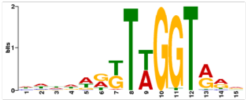 | 7.6 | GO:0009751<br>response to salicylic acid<br>GO:0009753<br>response to jasmonic acid<br>GO:0010224<br>response to UV-B<br>GO:0090379<br>secondary cell wall biogenesis involved in seed trichome differentiation |

**Figure S1. Selection of pathogen-inducible (PI) promoters based on transcriptomic analysis. a,** Venn diagram showing the overlap of significantly induced genes between different pathosystems. Pst, *Pseudomonas syringae pv tomato DC3000*; Pin, *Phytophthora infestans*; Xg, *Xanthomonas gardneri*; Xp, *Xanthomonas perforans*.**b,** The promoter sequences of 23 genes commonly induced by multiple pathogens were analyzed for significantly enriched transcriptional binding sites (q-value< 0.05) using the online TF binding site enrichment tool in PlantTFDB. Top binding sites related to stress response are listed.

a

Pst∆HopQ1 (top left), Xg∆XopQ (bottom left), Xe∆XopQ (top right) and $MgCl_2$ (bottom right) 6hpi

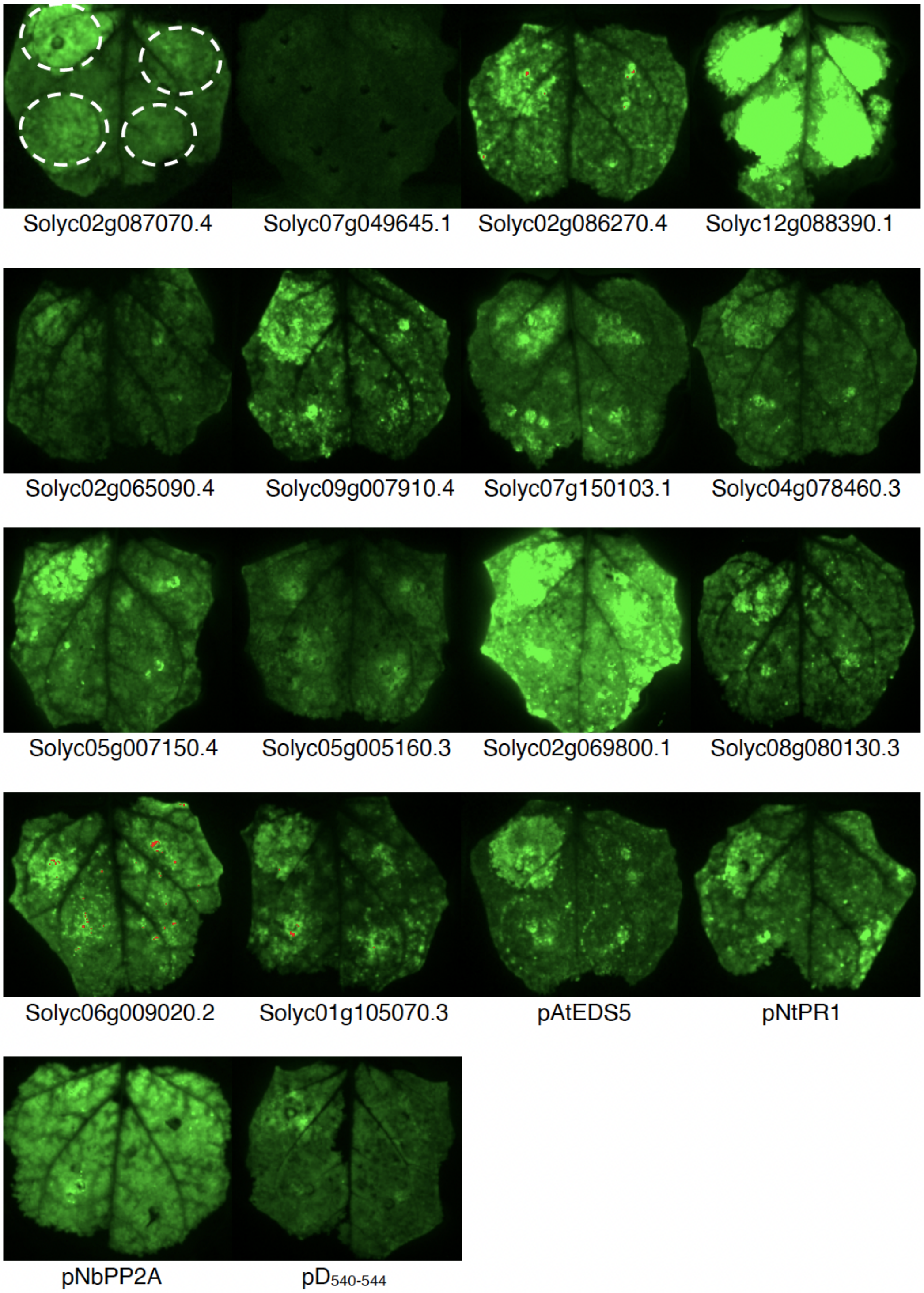


b

PstΔHopQ1, XgΔXopQ, XeΔXopQ and $MgCl_2$ 24hpi

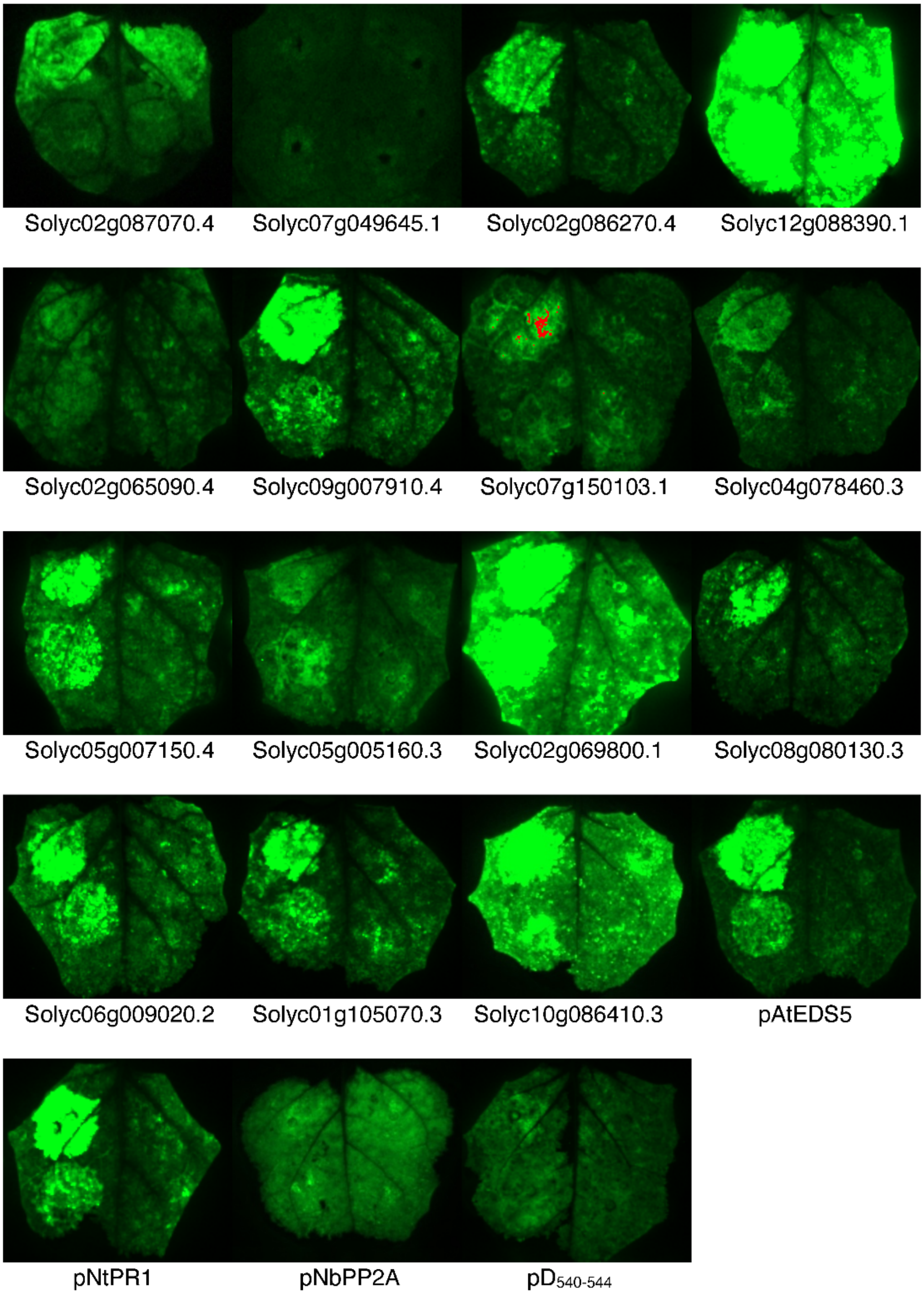


c

*P. capsici* 40hpi

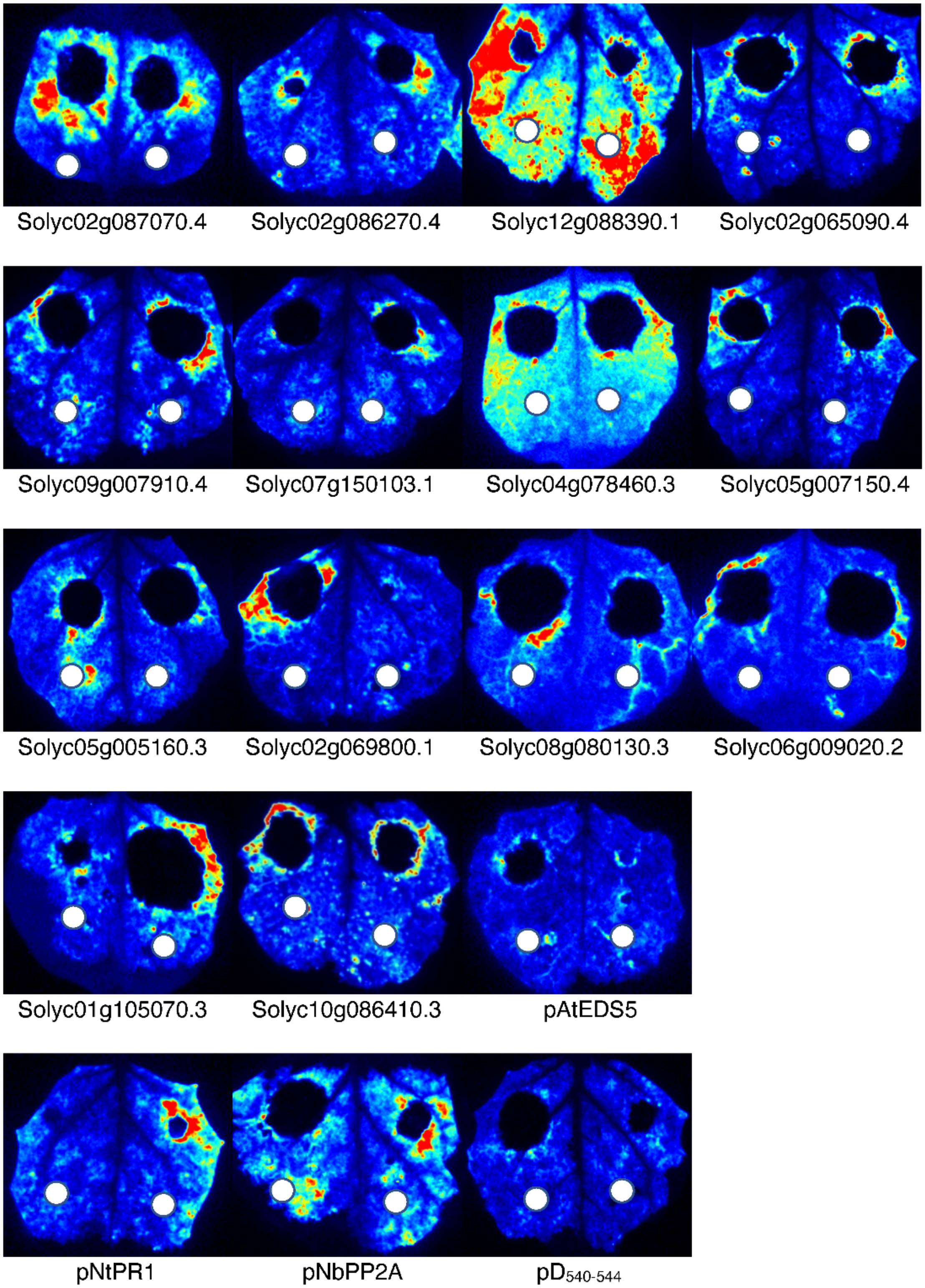

**Figure S2. Photos of *Nicotiana benthamiana* leaves transiently expressed with PI promoters regulating the luciferase reporter gene. a,b,** Response of PI promoters to bacterial pathogens 6 hours and 24 hours post infiltration. Top left, PstΔHopQ1; top right, XeΔXopQ; bottom left, XgΔXopQ; bottom right, $MgCl_2$ (mock). **c,** Response of PI promoters to *Phytophthora capsici* 40 hours post inoculation. White circles in the photos indicate sterile agar plugs that serve as a mock control.

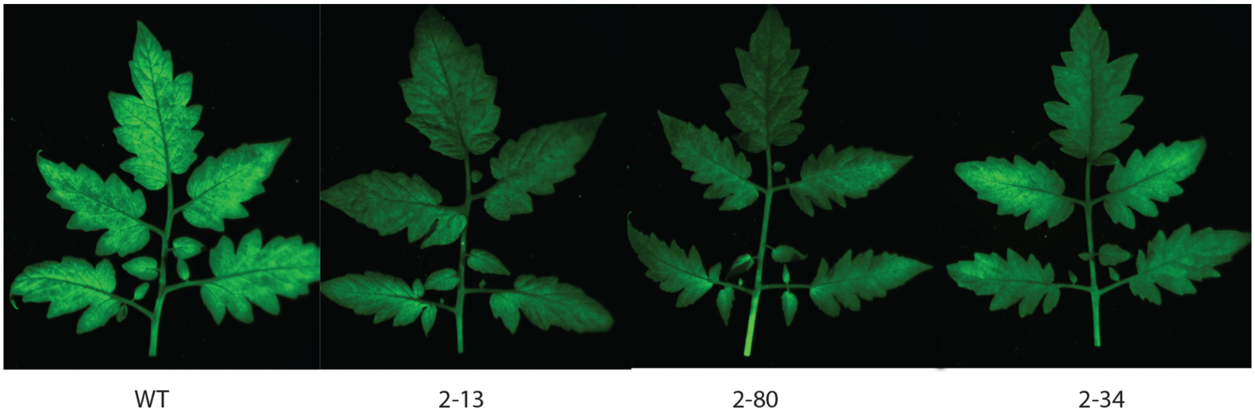


**Figure S3. Representative images of transgenic tomatoes infected with GFP-labeled *Pseudomonas syringae pv tomato DC3000*.** Phenotypes were imaged using 4 days post inoculation. Images were taken with a ChemiDoc MP imaging system (Biorad, USA) using excitation/emission wavelength for GFP. 2-13, 2-80 and 2-34 are independent transgenic events with *Solyc09g007910.4*::*Sr50.*

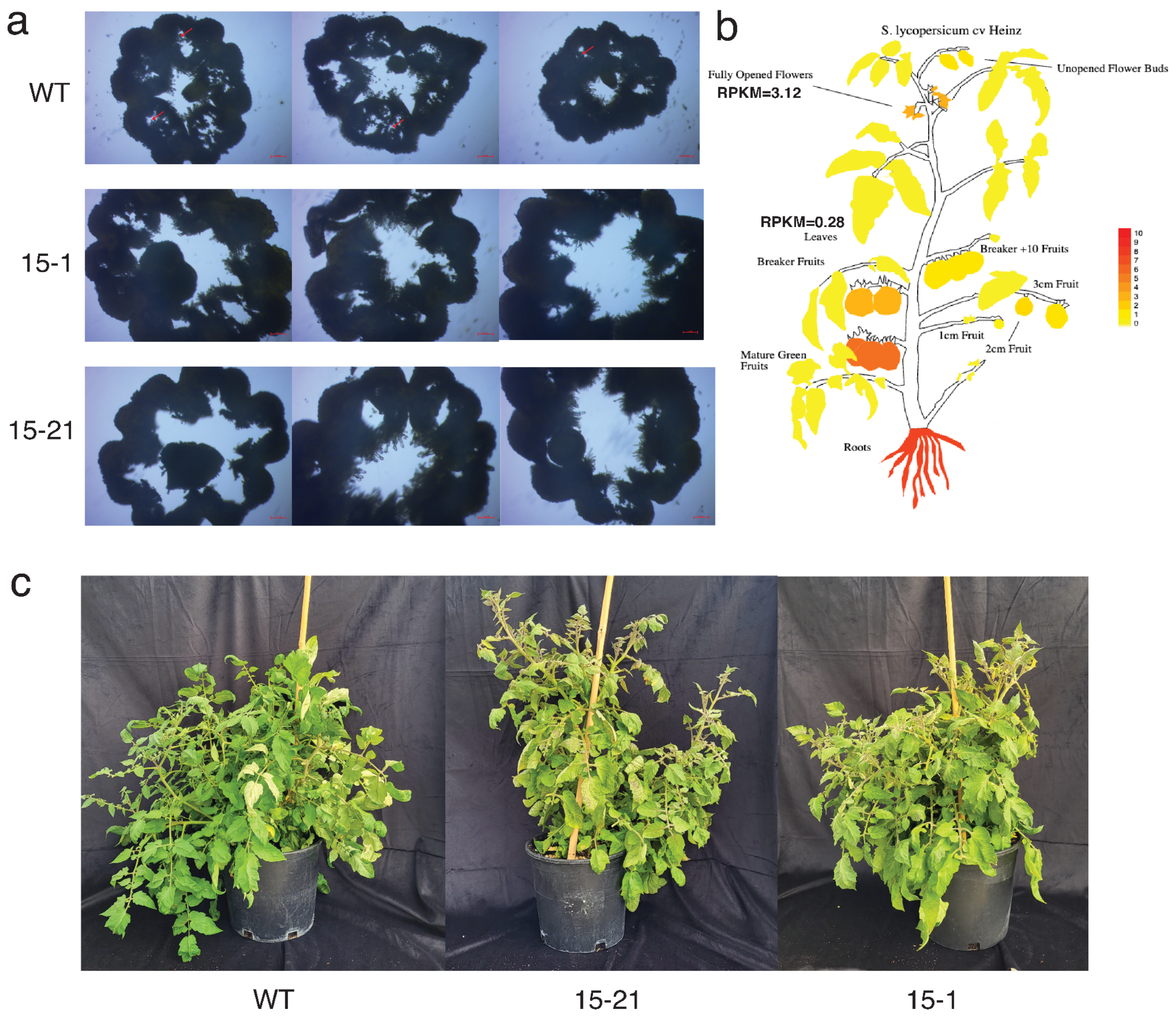


**Figure S4. Transgenic tomatoes expressing Sr33$^{E864V_V840M}$ regulated by promoter *Solyc02g086270.4* impairs reproduction but normal vegetative growth**. **a,** Cross-sections of fused stamens showing less pollen production in transgenics. **b,** Expression pattern of Solyc02g086270.4 in different tissues of WT tomato plants. The expression pattern and image was generated using the Plant eFP https://bar.utoronto.ca/eplant_tomato/. **c,** Transgenic plants showed similar vegetative growth as WT. Photos were taken 3 months after cutting.

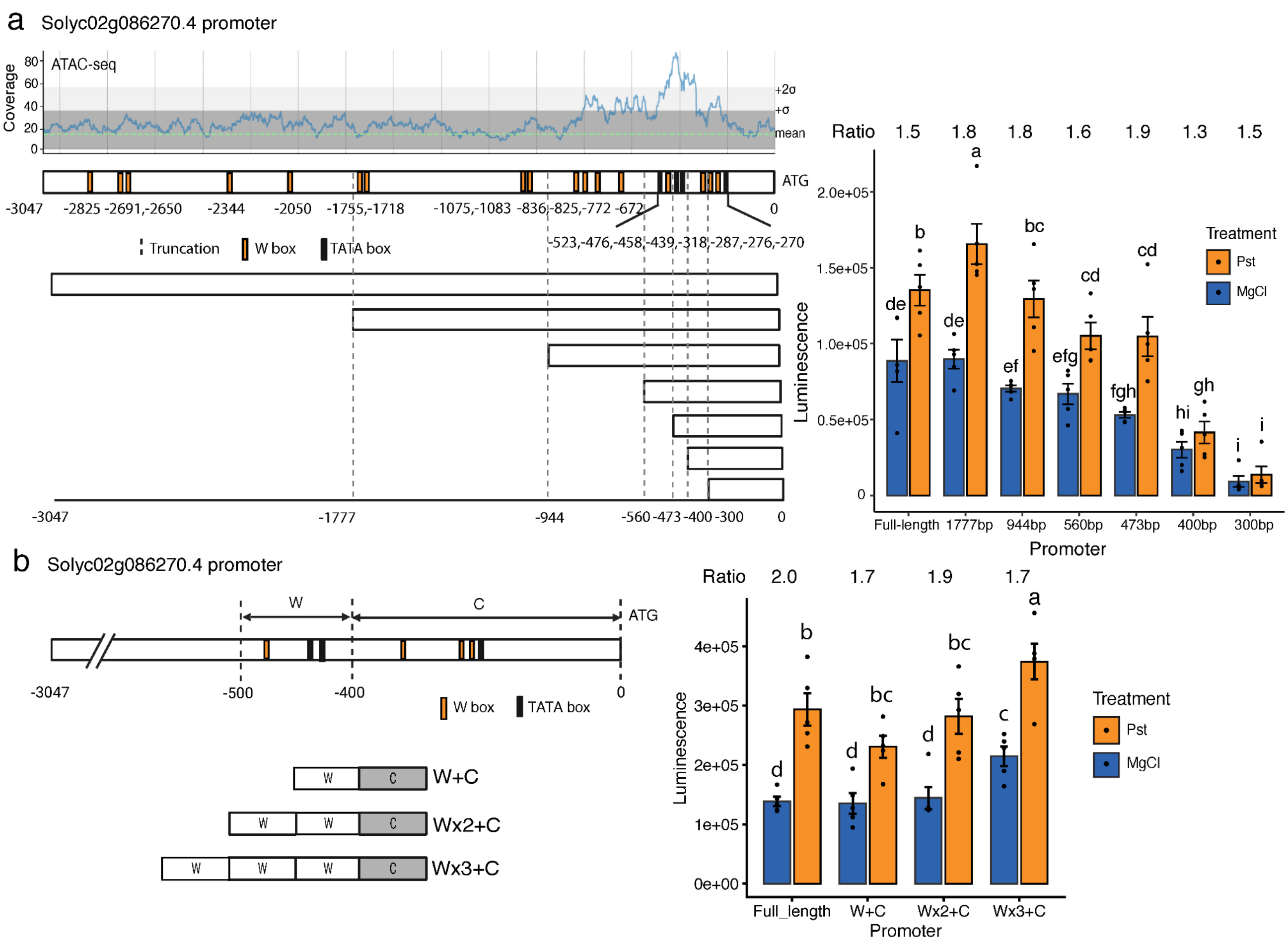


**Figure S5. Promoter engineering of Solyc02g086270.4 to modulate basal expression**. **a,** Identification of pathogen-inducible sequences in promoter by 5' end serial truncation. The top left shows the ATAC-seq read coverage plot of the promoter, with data from (Hendelman et al. 2021). The green dashed line indicates the genome-wide mean read coverage; the dark gray lines represent the mean ± 1 SD; and the light gray lines represent the mean ± 2 SD. The bottom left panel shows a schematic of the full-length promoter and truncation constructs. The bar plot on the right shows luciferase activity measured in *N. benthamiana* at 1 day post mock or PstΔHopQ1 infiltration. Statistical significance was assessed using two-way ANOVA followed by pairwise comparisons (n = 4-5). Different letters indicate significant differences ($p < 0.05$) after Benjamini-Hochberg multiple testing correction. **b,** Synthetic promoters with different repeat numbers of the pathogen-inducible sequence.

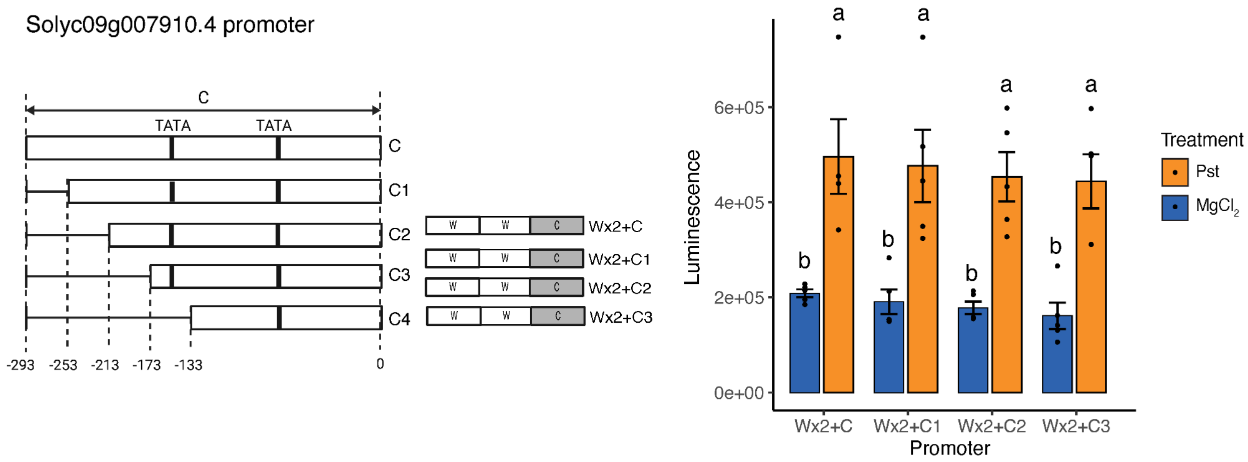


**Figure S6. Using different core promoter variants in the synthetic promoter did not show obvious gradient expressions**. The synthetic promoter only contains two copies of the inducible sequence. The bar plot on the right shows luciferase activity measured in *N. benthamiana* at 1 day post mock or PstΔHopQ1 infiltration. Statistical significance was assessed using two-way ANOVA followed by pairwise comparisons (n = 4-5). Different letters indicate significant differences ($p < 0.05$) after Benjamini-Hochberg multiple testing correction.
